# *In Silico* Toxicity Assessment of Organophosphates: A DFT and Molecular Docking Study on Their Interaction with Human Serum Albumin (HSA)

**DOI:** 10.1101/2024.12.15.628551

**Authors:** Tanya Singh, Nisha Shankhwar, Neeta Raj Sharma, Anil Kumar, Awadhesh Kumar Verma

**Affiliations:** School of Bioengineering and Biosciences, Lovely Professional University, Punjab, 144411, India+; Special Centre for Nanoscience, Jawaharlal Nehru University, New Delhi, 110067, India; Gene Regulatory Laboratory, National Institute of Immunology, New Delhi, 110067, India

**Keywords:** *In silico* Toxicity Assessment, Organophosphates, DFT, Molecular Docking, Human Serum Albumin

## Abstract

This study presents an *in silico* analysis of the toxicity of Organophosphates (OPs) through their interaction with Human Serum Albumin (HSA) protein using density functional theory (DFT) and molecular docking approaches. Organophosphates, known for their widespread use as pesticides, have raised significant concerns due to their potential toxicological effects. To investigate the molecular mechanisms underlying OP toxicity, we conducted DFT calculations to determine the electronic properties and reactivity of selective OP compounds. Molecular docking simulations were performed to explore the binding affinity, interaction sites, and conformational changes of HSA upon exposure to OPs. The DFT analysis revealed key electronic descriptors, such as HOMO-LUMO gap and electrostatic potential, that indicate high reactivity of OPs with biological molecules. Docking results showed strong binding affinities between OPs and HSA, particularly at sites involved in metabolite and drug transport, suggesting potential interference with the protein’s native function. The interaction of OPs with HSA was further supported by molecular dynamics simulations, which confirmed the stability of the OP-HSA complex over time. These findings provide critical insights into the molecular basis of organophosphate toxicity, emphasizing the importance of their interaction with HSA. The combined DFT and molecular docking approach offers a valuable framework for predicting the toxicological behavior of OPs and lays the foundation for further *in vitro* and *in vivo* studies.

**Graphical Abstract:** 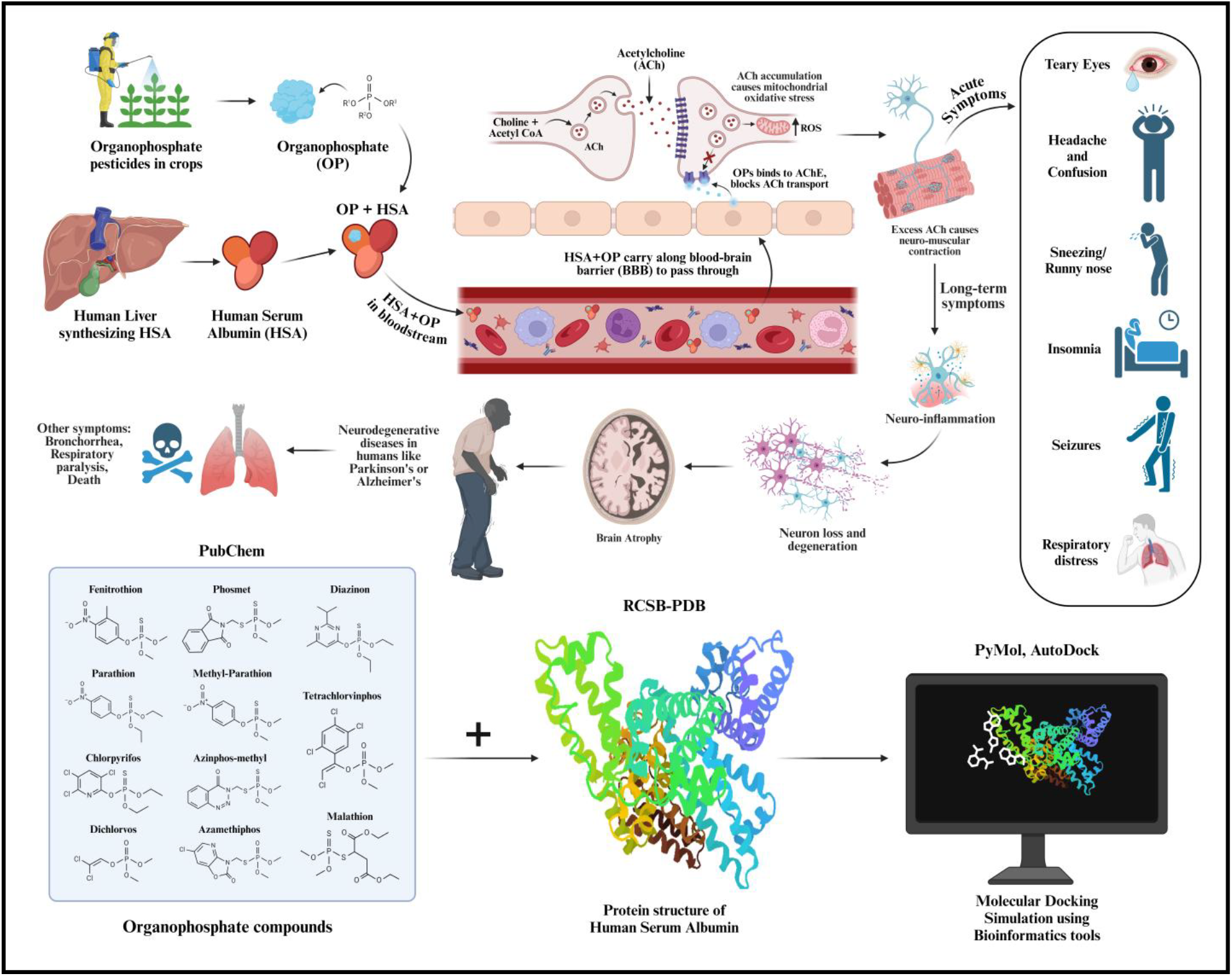

## 1. Introduction

Organophosphates (OPs) comprises a class of organo-phosphorus compounds, also known as the esters of phosphoric acid and alcohol. It’s main application has been observed in making pesticides, insecticides, flame retardants, engine oil additives for polymer and plastic durability and manufacture of nerve agents for war conflicts. Toxicity regarding organophosphates can harm humans through accidental exposure to pesticides and insecticides or as an occupational hazard while production, through terrorist activities or chemical warfare, or through intentional poisoning [1]; it also has environmental impacts where due to agricultural overexploitation, the residues of OPs can seep through soil causing leaching and accumulation, thereby contaminating groundwater and unintentionally harming the terrestrial and aquatic food chains [2]. This further can cause acute exposure and toxicity to insects, plants, animals and humans as well. Organophosphates are generally known for inhibition of enzyme Acetylcholine-esterase in land or marine organisms, and their development as pesticides is targeted towards pests/insects who harm the cash crops [3]. The inhibition can cause respiratory, nervous, hepatic (liver related) and reproductive malformations. Among plants, OPs cause disarray in the growth promoting mechanism by inhibiting function of key plant growth enzymes, transcuticular diffusion and permeability for essential nutrients in plant cells [4]. In humans, OPs as bioactive compounds can disrupt the functioning of acetylcholine-esterase enzyme (AChE) in the nervous system by making a covalency with its active site, which is found in the post-synaptic junctions between the nerve cells and muscle cells [5]. Normally, during motor nerve stimulation, the neurotransmitter compound acetylcholine is released for transmitting neural impulses to organs and muscles. Then by the process of hydrolysis the enzyme breaks down acetylcholine to relax the organ or muscle. But when OPs get bound, they disturb the functioning of AChE which causes accumulation of acetylcholine and leading to non-stop transmission of nerve impulses and contraction of muscle cells [6]. This also affects the organs and glands through this continual cycle of impulses and contraction resulting in immediate symptoms such as tearing of eyes, uncontrolled drooling or foaming, and excess nose mucus production causing rhinorrhea. In case of acute harm caused by OPs, symptoms may vary from anxiety, confusion, headaches, drowsiness, seizures, insomnia, memory loss, to circulatory or respiratory depression [7]. Chronic organophosphate toxicity could lead to the patient being unresponsive, having eye miosis or pinpoint pupils, involuntary muscle twitching, diaphoresis or excessive sweating and rapid uremia. In the cases of fatality, the frequent cause is respiratory failure occurring due to broncho-constriction or tightening of smooth muscles which constricts the lungs’ airways, bronchorrhea or production of watery sputum by more than 100 ml by the patient, respiratory depression and acute muscle paralysis in the respiratory system [8]. Long-term effects of acute poisoning have also been observed to occur among the survivors of nerve agents. OPs could meddle with mitochondrial functioning among nerve cells causing oxidative stress, leading to neurogenerative diseases like Alzheimer’s disease, Parkinson’s disease, ALS (Amyotrophic Lateral Sclerosis), and neurotoxic diseases like OPIDN (Organophosphate Induced Delayed Neuropathy), COPIND (Chronic Organophosphate Induced Neuropsychiatric Disorder) and Cholinergic syndrome [8],[9].

Human Serum Albumin (HSA) is a heart-shaped globular protein consisting of six repetitive helical sub-domains, considered the principal element in the human serum concentration. It is synthesized by hepatocytes, constituting about 40% of the daily protein synthesis by liver cells [10], and helps in regulating plasma osmotic pressure in blood and transports ligands such as bilirubin, fatty acids, warfarin, ibuprofen and various drugs at the right active sites in human body [11]. The interaction between OPs and HSA has shown to be associated with toxico-kinetics (process explaining ADMET) and toxico-dynamics (signs and symptoms a toxic agent causes while interacting with a biological target) in human body, depending on the bioavailability [12]. Many studies regarding OPs and HSA interaction have been done on the basis of clinical studies and computational analysis. One such study referred to estimating whether the concentration of serum albumin was associated with mortality in patients suffering from hypoalbuminemia at presentation [13]. This study was conducted upon 217 patients exposed to organophosphate (OP) poisoning, and the disease at presentation, whose serum albumin is considered less than 3.5 g/dL was identified in 18.4% of the patients poisoned with OP [13]. Another investigated the interactions of OPs such as Parathion-methyl and Malathion with HSA, where through Solid-phase micro-extraction (SPME) methods the value of G (the thermodynamic constant) obtained was found to be almost similar to the molecular docking values, and it was deduced that the mode of interaction between HSA and Parathion-methyl, HSA and Malathion were bonded chiefly by hydrogen bonds, pi-to-pi stacking and hydrophobic effects [14]. Another study was related to the interaction of Chlorpyrifos with HSA and the effects of Diazinon on HSA structural changes. It was observed that effects caused by Diazinon were through interaction with the amino acid TRP 214 residue present in HSA, and binding of the ligand with TRP 214 was confirmed by molecular docking. Further, the hydrophobic and electrostatic nature of binding pattern was analyzed in Diazinon interaction with HSA [15]. It was also observed that Parathion and Paraxon interact with one kind of HSA binding site, where the secondary structures of the ligands changed when interacting with the binding site of the Chlorpyrifos on HSA receptor TRP 214, which could have been stabilized by hydrophobic and electrostatic interactive forces [15].

Aside from how toxicity caused by OPs can have adverse effects, there are also studies done for reducing the intoxication in environmental and human life aspect. One such study had researched about how by the use of bio-scavengers as catalytic components can help minimize effects of nerve agents in humans, by targeting their isomers and through the process of hydrolyzation of compounds help in counteracting effects of OPs, although this study is at experimental stage and has been done on animal subjects at present [16]. Other study focused on bioremediation through microbial decontamination for removing hazardous chemicals from the soil for sustainable agriculture. While discussing about what detrimental effects can OP pesticides cause to the human body, the study also reflects on the necessity of screening probiotic strains which are resistant to pesticides along with conducting bioremediation studies through in-vivo and in-vitro probiotic models [17].

Our focus for this study would be on comparing and analyzing the interactions of different components of organophosphate pesticides with HSA at molecular level and to determine if their interactions differ in terms of covalent or non-covalent bonding, interaction with the binding site, along with emphasis on toxicity of these components on human health [18].

## 2. Methodology

### 2.1 Modeling and optimization of organophosphate pesticides as ligands

The three-dimensional (3D) initial modeling of 12 Organophosphates (Azamethiphos, Azinphos-methyl, Chlorpyrifos, Diazinon, Dichlorvos, Ethion, Fenitrothion, Malathion, Parathion, Methyl-parathion, Phosmet and Tetrachlorvinphos) commonly known as organophosphorus pesticides, was done through MarvinSketch software [19]. The molecular structures were initially refined in both 2D and 3D forms, then the geometries of the OP pesticide components were optimized to their transition states. Intermediate energy minimizations were performed through ChemDraw software for progressive refinement of molecular geometries [20]. Further optimization was conducted using Density Functional Theory (DFT) with Gaussian09 software package. The DFT calculations employed the RB3LYP function in combination with the 6-311G(d,p) basis set, offering a high level of computational accuracy for electronic structure calculations [21],[22]. The optimized molecular geometries were then used in molecular docking studies to investigate the interactions between the organophosphate compounds and human serum albumin (HSA), a key transport protein. The docking analysis provided insights into the binding affinities and interaction patterns between the ligands and HSA [23].

### 2.2 Preparation of HSA protein as Receptor molecule

The 3D structure of HSA was retrieved from the RCSB Protein Data Bank (PDB ID: 7VR0), derived from Homo sapiens. The structure has a resolution of 1.98 Å and consists of 585 amino acid residues, without any reported mutations. Before conducting docking studies, the HSA protein structure underwent cleaning processes by AutoDockTools 4.2 [24]. This involved the removal of all crystallographic water molecules, ligands, and any other crystallizing agents that might interfere with the docking simulation. Then, polar hydrogens were added to the protein structure to ensure proper hydrogen bond interactions during docking [25]. Kollman charges were assigned to the atoms, and Gasteiger charges were computed to accurately represent the electrostatic properties of the protein. The atomic types were then defined as AutoDock4 (AD4) types, ensuring compatibility with the AutoDock docking algorithm [26].

### 2.3 Molecular Docking of Organophosphates with HSA molecules

Molecular docking in rigid form was performed using the optimized OP pesticide ligand molecules and the HSA protein structure. The docking simulations were carried out through AutoDock tools version 4.2, which employed the use of Genetic algorithm (GA) simulation approach [27]. The GA was configured with a population size of 300 and was run for 100 iterations to ensure a comprehensive search of the ligand conformations and binding poses. All other docking parameters were made to maintain their default values to standardize the experiment [28]. The output docked files were saved using the Lamarckian Genetic Algorithm (LGA) method, which combines the genetic algorithm’s global search capabilities with local optimization strategies for higher accuracy in docking predictions [29]. The first 10 conformations of the HSA protein-OP ligand complexes were selected based on their most favorable negative binding energies (ΔG) and low root mean square deviation (RMSD) values, indicating strong binding affinities and stability of the docked conformations [30]. Post-docking analysis was performed using PyMOL and Discovery Studio software. These tools were employed to visualize and study the molecular interactions between Organophosphates and HSA, as well as to analyze binding energies, interaction sites, and conformational changes in the complexes [31].

## 3. Results and Discussion

### 3.1 Modeling and docking studies on Organohosphate-HSA Complex using *in silico* approach

Figure 1 shows the ground state structure of OP pesticide (example model here taken as Azinphos-methyl) generated using ChemDraw software, as a ball and stick model. The grey colored balls represents the carbon atoms, white colored ones as hydrogen atoms, red colored as oxygen atoms, blue colored as nitrogen atoms, dark yellow colored as sulphur atoms and pink colored as phosphorus atom in configuration of Azinphos-methyl. Entire structure has two hexagonal rings attached to each other through nitrogen, carbon and oxygen atoms, further which were attached to central phosphorus atom via carbon and sulphur atoms.

**Figure 1:**
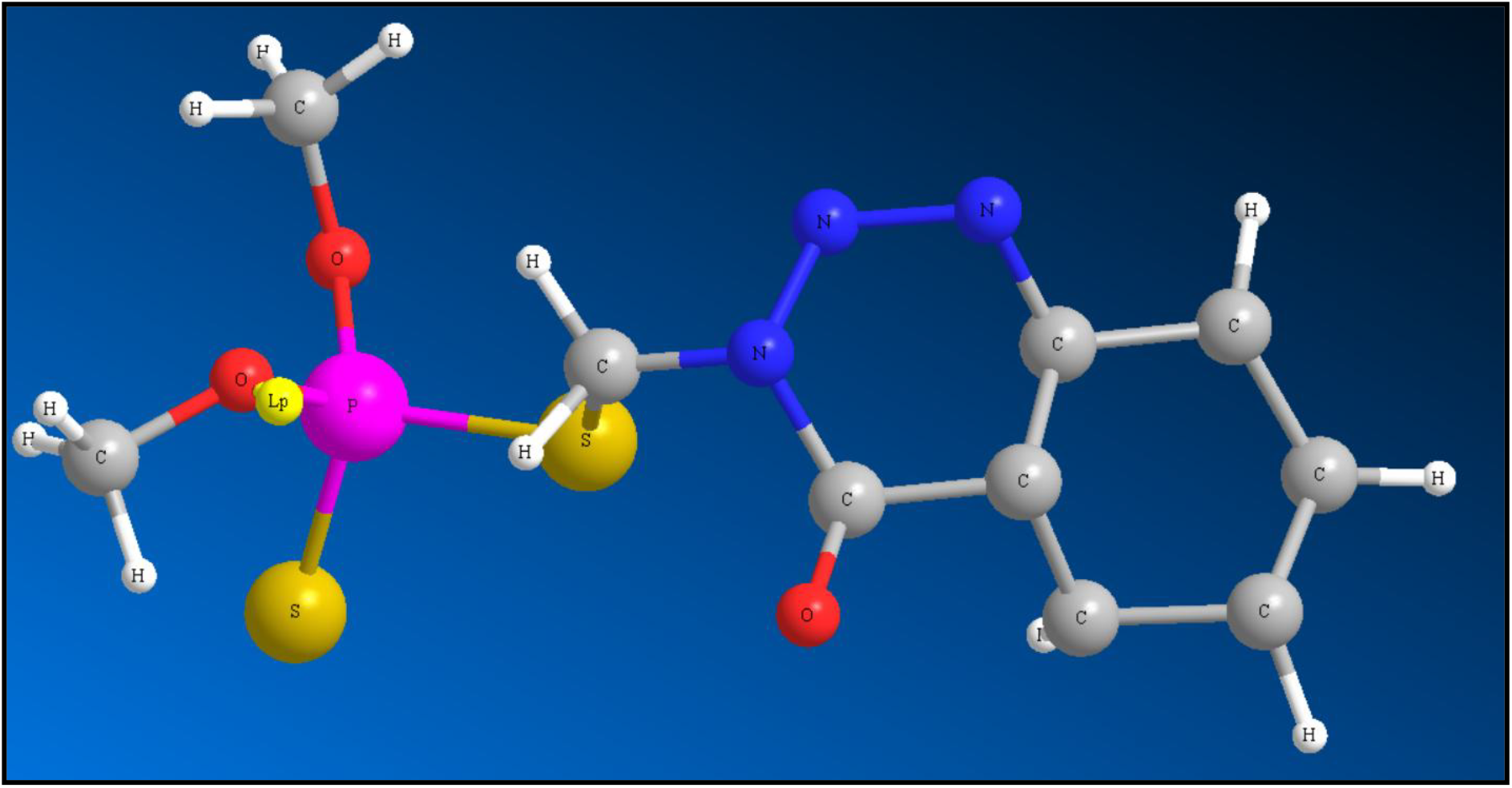
3D ground state configuration generated for Azinphos-methyl compound using ChemDraw software.

Optimization and energy minimization of the OP compounds was done using Gaussian09 software after being generated through ChemDraw. The graph obtained showed how much of optimized step numbers were taken to achieve total energy minimisation for the OP compound Azinphos-methyl. Then the optimization of the compound was done through the calculation of bond length, bond angles and the types of bond angle for each atom of Azinphos-methyl molecule.

The detailed information regarding bond length, bond angle and dihedral angle for other OP pesticide compounds including their energy minimization and optimization step number has been shared in supplementary data. After optimisation, the bond lengths in OP molecular structure showed no change while bond angle showed some slight changes.

### 3.2 Molecular docking of OP pesticides with HSA protein

Here, computational approaches like molecular docking were also analysed using AutoDockTools software, so as to give insights for the molecular interaction between OPs and HSA. After obtaining results from the molecular docking, it was observed that all the OP pesticide compounds showed negative biniding affinities with HSA protein on the basis of docking score, computationally proving our theory that pesticides do have a strong binding property with human albumin proteins which cause biological toxic effects. The docking scores have been compiled for 12 OP pesticide compounds in Table 1, and their docking scores are plotted as a graph in Figure 2.

**Table 1:** Affinity-based selection of organophosphates – HSA complex: analysis of docking scores for different pesticides.

| OP pesticides | Docking score against HSA<br>(kcal/mol) |
| --- | --- |
| **Azinphos-methyl** | **-5.75** |
| **Fenitrothion** | -5.62 |
| **Tetrachlorvinphos** | -5.17 |
| **Phosmet** | -5.16 |
| **Methyl-parathion** | -5.08 |
| **Parathion** | -4.98 |
| **Diazinon** | -4.97 |
| **Azamethiphos** | -4.44 |
| Dichlorvos | -3.68 |
| Chlorpyrifos | -3.66 |
| Ethion | -2.81 |
| Malathion | -1.90 |

**Figure 2:**
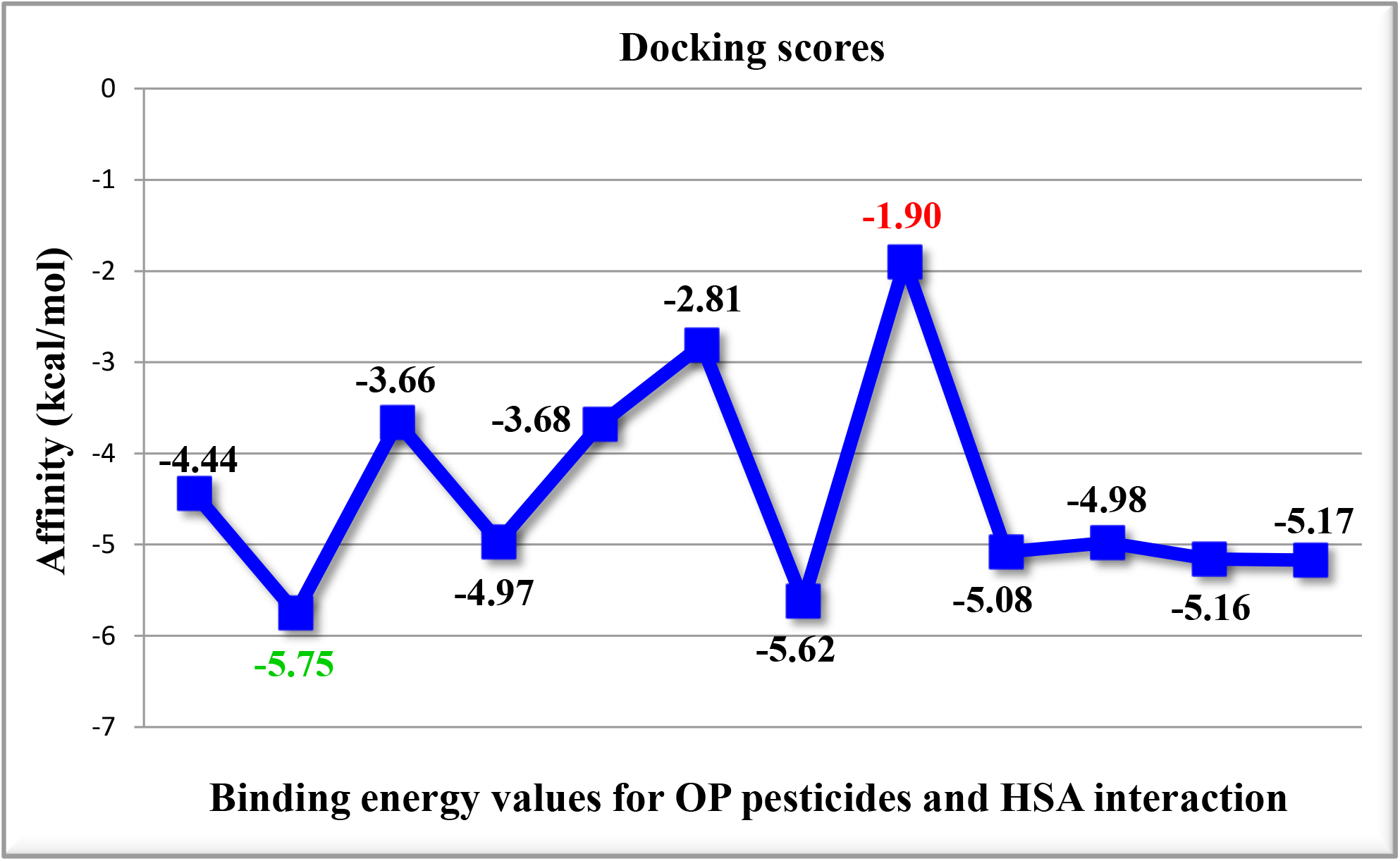
Graphical plot for docking scores of Organophosphate pesticides (ligand) with HSA (protein). More negative scores indicate more favorable binding affinity.

Table 2 shows the compiled rank wise different confirmations of 12 OP compounds and HSA complex. For Azinphos-methyl, the best binding energy came out to be -5.75 Kcal/mol with cluster RMSD as 0.00 and reference RMSD to be 18.26. Similarly, for the rest 11 compounds, their first rank results have been tabulated below with respective RMSD values.

**Table 2:** Confirmation of OP-HSA complex by rank with respective binding energy and RMSD value.

| OP pesticides | Rank | Run | Binding Energy | Cluster RMSD | Reference RMSD |
| --- | --- | --- | --- | --- | --- |
| Azamethiphos | 1 | 9 | -4.44 | 0 | 18.54 |
| **Azinphos-methyl** | **1** | **8** | **-5.75** | **0** | **18.26** |
| Chlorpyrifos | 1 | 3 | -3.66 | 0 | 11.16 |
| Diazinon | 1 | 7 | -4.97 | 0 | 11.87 |
| Dichlorvos | 1 | 1 | -3.68 | 0 | 18.76 |
| Ethion | 1 | 7 | -2.81 | 0 | 26.78 |
| Fenitrothion | 1 | 8 | -5.62 | 0 | 18.54 |
| Malathion | 1 | 3 | -1.9 | 0 | 43.33 |
| Methyl-parathion | 1 | 9 | -5.08 | 0 | 45.04 |
| Parathion | 1 | 9 | -4.98 | 0 | 44.39 |
| Phosmet | 1 | 1 | -5.16 | 0 | 18.18 |
| Tetrachlorvinphos | 1 | 6 | -5.17 | 0 | 17.43 |

Figures 4(a) and 4(b) show the 2D and 3D docked structure of OP Azinphos-methyl with HSA compiled using Discovery Studio software where some parts of the protein’s amino acid residues showed involvement in non-covalent interaction with Azinphos-methyl, deducing that the ligand compound is bound to the protein’s active site. For the van der Waals interaction, it was observed through amino acids LEU 189, LYS 190, ILE 142, LEU 115, PRO 118 and GLU 141. TYR 161, LYS 137 and PHE 134 showed pi-donor hydrogen bonding and carbon-hydrogen bonding with the ligand molecule, respectively. MET 123 demonstrated pi-sulphur bonding, while TYR 138 showed pi-pi T-shaped interaction. action. ARG 117, 186 and LEU 182 had pi-alkyl interaction with the benzene rings of the ligand structure. It was also observed that TYR 161 was involved majorly in pi-hydrogen, carbon-hydrogen, pi-sulphur and pi-pi T-shaped bonding with the Azinphos-methyl structure, indicating that it was the main amino acid of the HSA’s active site where the OP compound Azinphos-methyl strongly bound to.

**Figure 4(a):**
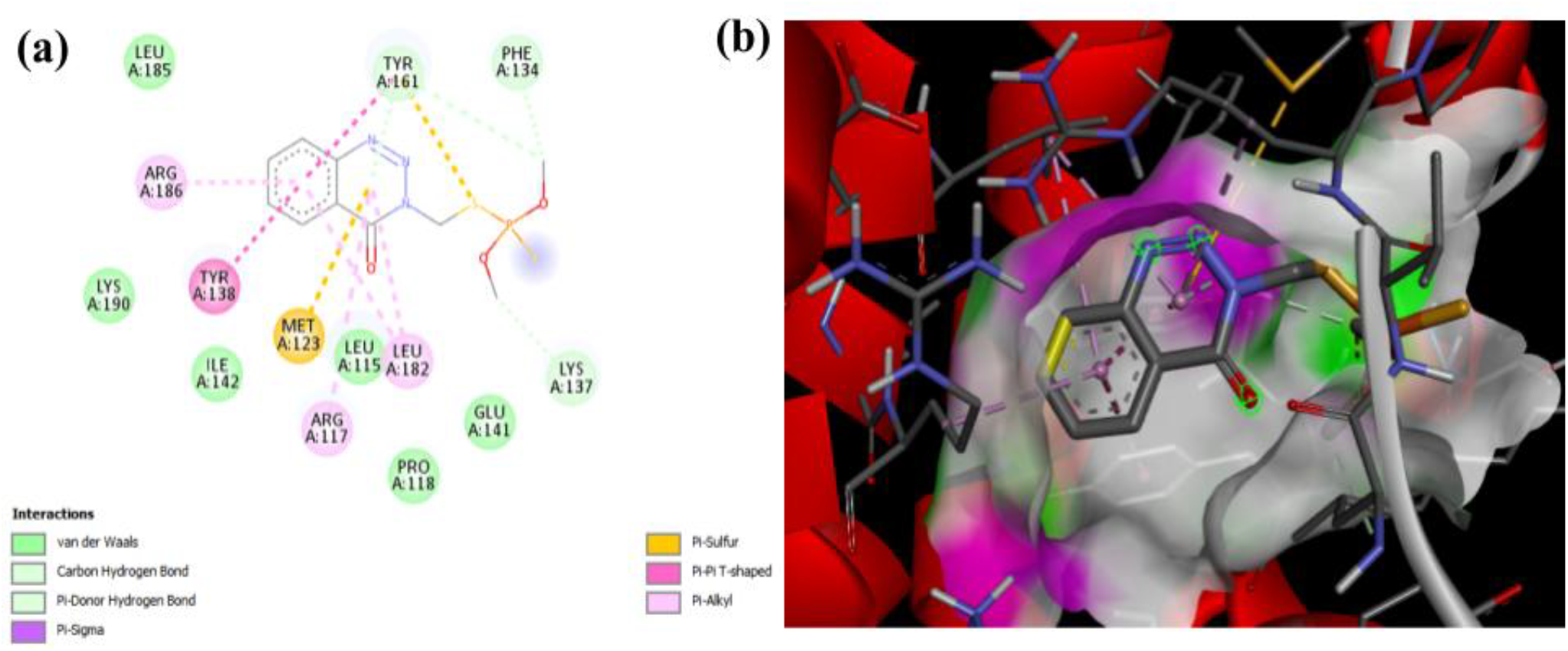
2D representation of non-covalent bonding between Azinphos-methyl and HSA complex; 4(b): 3D interaction of Azinphos-methyl with HSA modeled using PyMOL and Discovery Studio software.

From the docking study and deduction of binding affinity, it is clear that the OP pesticide Azinphos-methyl shows great binding affinity with HSA protein compared to other compounds and can be treated as a potential OP compound causing toxicity in human beings.

### 3.3 Visualization and analysis of Organophsphate-HSA complex formation using LigPlot plus

Softwares like LigPlot Plus helped further analyze the complex, highlighting specific interactions of OPs with HSA protein, most likely hydrophobic interactions and hydrogen bonding contributing to the complex stability. The docking complex obtained through LigPlot plus revealed a network of interactions that can be seen in Fig 5(a) for HSA and Azinphos-methyl. Predominant among these forces are hydrophobic bonds and non-covalent bonding among the amino acids of the protein, where a peculiar hydrogen bond of bond length 3.27 Å was observed between ARG 257 amino acid of HSA and an oxygen molecule attached to central phosphorus atom of Azinphos-methyl. The potential presence of salt bridges with ligand bonds further suggests a role for electrostatic attractions in complex formation. In Fig 5(b), similar observations can be seen for Malathion interaction with HSA, which showed least binding affinity, but that complex is stabilized only by hydrophobic interactions, ligand bonds and external bonds.

**Figure 5(a):**
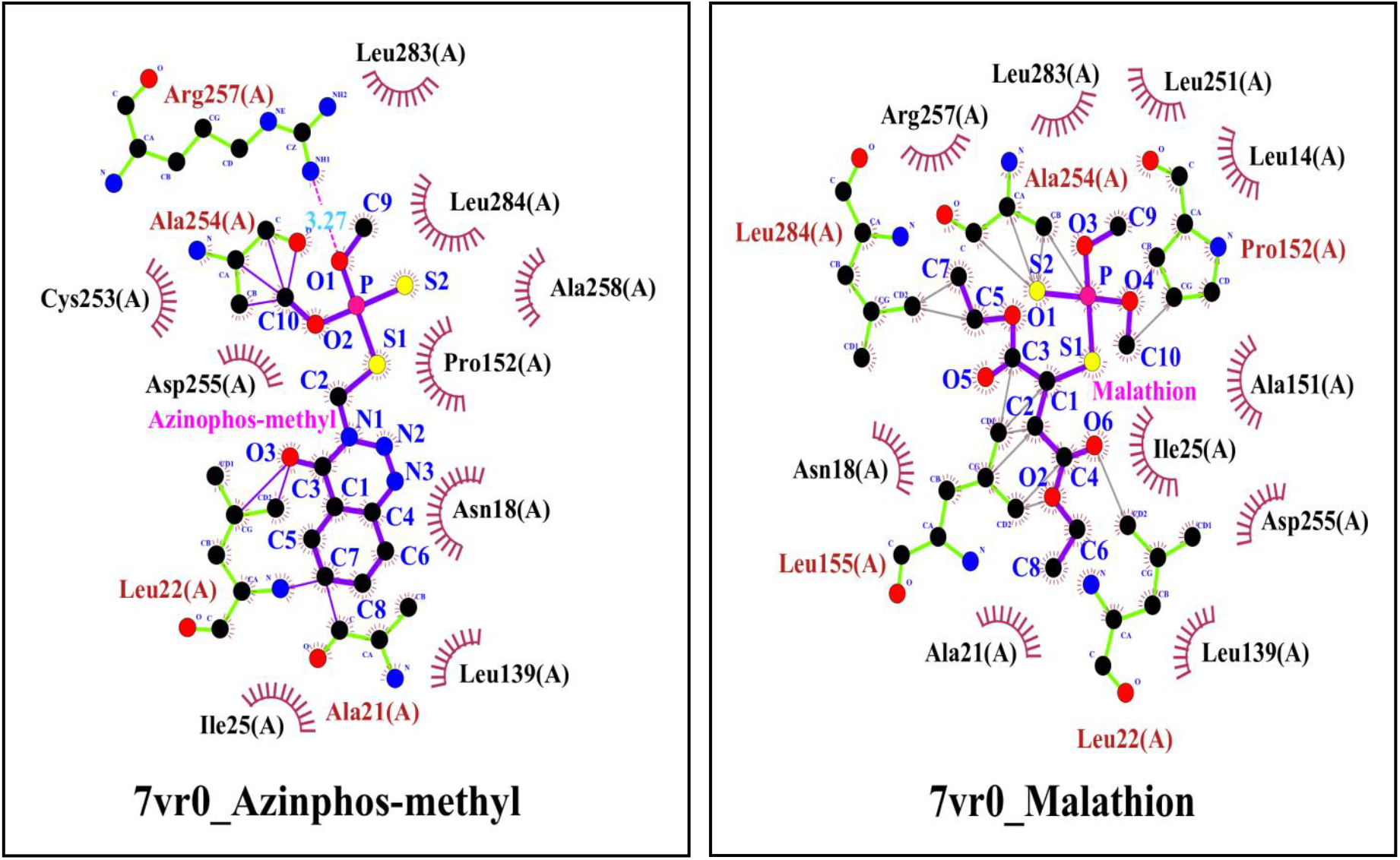
Snapshot showing the interaction between HSA and pesticide molecule Azinphos-methyl using LigPlot plus software; 5(b): Snapshot showing the interaction between HSA and pesticide molecule Malathion using LigPlot plus software.

### 3.4 Toxicity assessment and analysis of Organophosphate pesticide compounds

To analyse the toxicodynamics of these pesticide compounds of organophosphorus nature, it was assessed using Swiss-ADME software and ProTox 3.0 tools. Using the SMILES ID of these compounds, the toxicity prediction chart was generated for each of them through these tools and it was deduced that most of these compounds fall under high level of toxicity class, indicating that they are highly toxic in nature and are bound to cause toxic effects to different organs. Table 3 shows the compiled toxicity analysis for all 12 OP pesticides in the form of ADMET table, where for the compound Azinphos-methyl, its predicted toxicity class was 1 meaning it has high toxicogenicity.

**Table 3:** ADMET Table for different OP pesticides obtained from Swiss-ADME.

| OP pesticides | Compound ID | Molecular weight (g/mol) | H-bond Acceptor | H-bond donor | Log P | Molecular refractivity | Predicted Toxicity class |
| --- | --- | --- | --- | --- | --- | --- | --- |
| Azamethiphos | 71482 | 324.68 | 6 | 0 | 1.3 | 72.58 | 4 |
| Azinphos-methyl | 2268 | 317.32 | 5 | 0 | 2.16 | 80.00 | 1 |
| Chlorpyrifos | 2730 | 350.59 | 4 | 0 | 4.12 | 77.92 | 3 |
| Diazinon | 3017 | 304.35 | 5 | 0 | 3.23 | 80.23 | 2 |
| Dichlorvos | 3039 | 220.98 | 4 | 0 | 1.6 | 42.38 | 2 |
| Ethion | 3286 | 384.48 | 4 | 0 | 4.07 | 96.04 | 2 |
| Fenitrothion | 31200 | 277.23 | 5 | 0 | 2.05 | 69.27 | 3 |
| Malathion | 4004 | 330.36 | 6 | 0 | 2.14 | 78.08 | 3 |
| Methyl-parathion | 4130 | 291.26 | 5 | 0 | 2.42 | 64.30 | 2 |
| Parathion | 991 | 263.21 | 5 | 0 | 1.71 | 73.92 | 1 |
| Phosmet | 12901 | 317.32 | 4 | 0 | 2.14 | 81.66 | 2 |
| Tetrachlorvinphos | 5284462 | 365.96 | 4 | 0 | 3.89 | 77.89 | 4 |

Table 4 shows the compiled analytical results for 12 compounds describing their active or inactive role in causing neuro-, hepato-, respiratory, nephro- and cardiotoxicity, along with critical factors like gastrointestinal (GI) absorption and blood-brain barrier (BBB) permeability. Among these, the compound Azinphos-methyl showed high GI absorption and ‘Active’ status in causing neurotoxicity.

**Table 4:** Organ Toxicity chart for OP pesticides obtained through ProTox-3.0.

| OP pesticides | GI absorption | BBB permeable | Neuro-toxicity | Hepato-Toxicity | Respiratory toxicity | Nephro-toxicity | Cardio-toxicity |
| --- | --- | --- | --- | --- | --- | --- | --- |
| Azamethiphos | High | No | Active | Active | Active | Active | Inactive |
| Azinphos-methyl | High | No | Active | Inactive | Inactive | Inactive | Inactive |
| Chlorpyrifos | High | No | Inactive | Inactive | Inactive | Active | Inactive |
| Diazinon | High | No | Inactive | Inactive | Inactive | Inactive | Inactive |
| Dichlorvos | High | Yes | Inactive | Inactive | Active | Inactive | Inactive |
| Ethion | Low | No | Inactive | Inactive | Inactive | Inactive | Inactive |
| Fenitrothion | High | No | Inactive | Inactive | Inactive | Inactive | Active |
| Malathion | Low | No | Inactive | Inactive | Active | Active | Inactive |
| Methyl-parathion | High | No | Inactive | Inactive | Inactive | Inactive | Active |
| Parathion | High | No | Inactive | Inactive | Inactive | Inactive | Active |
| Phosmet | High | No | Inactive | Inactive | Active | Active | Inactive |
| Tetrachlorvin phos | High | Yes | Inactive | Inactive | Inactive | Inactive | Inactive |

## Conclusion

Through analysis for organophosphates and their predictive toxicogenicity, our study came to the conclusion that OP toxicity can abruptly influence human biological processes and cause the aforementioned diseases, and for that, computational analysis of interactions of different OP compounds with the human serum albumin protein was required to prove our research theory. The optimization of 12 OP pesticide compounds by modeling their chemical structures for this study demonstrated difference in the physical and conformational structures for each compound in terms of bond angle, bond length, minimization of energy for getting stable ligands for molecular docking, visualization of protein-ligand interaction and toxicity analysis. Molecular docking showed Azinphos-methyl to have best docking score among the rest of OP pesticides, giving us the impression that it can be considered a promising compound to be studied further for toxicity prediction. After 2D and 3D visualisation of the pesticide compounds through PyMol and Discovery Studio, the detailed conformations regarding the type of bonds and protein amino acids stabilizing the 12 OP pesticide-HSA complexes respectively, were identified and picturized through LigPlot software. These *in silico* results could help in intervention of efficacy of OPs when bound with transport proteins like HSA and being carried to their targets like AChE, so that potential and effective antidotes can be developed to minimise damage to human organs caused by OPs. Further computational analysis would be required to study the interaction of HSA with other types of OPs like nerve agents, chemicals for medical use and industrial application chemicals. Therefore, through this study we not only got the biological insight for the action and interaction of organophosphate pesticides with essential human proteins like HSA, but also through computational modeling we were able to learn how such molecules can be made a target for drug development and toxicological evaluation, along with taking steps for minimising their overuse in agriculture so as to protect and conserve human health and our environment.

## Supporting information

Supplementary_files_OP_HSA

## Abbreviations

OP: Organophosphates
HSA: Human Serum Albumin
DFT: Density Functional Theory
HOMO-LUMO gap: Ratio of Highest Occupied Molecular Orbital to Lowest Unoccupied Molecular Orbital gap
AChE: Acetylcholinesterase
ALS: Lou Gehrig’s disease (Amyotrophic Lateral Sclerosis)
OPIDN: Organophosphate Induced Delayed Neuropathy
COPIND: Chronic Organophosphate Induced Neuropsychiatric Disorder
SPME: Solid-Phase Micro-Extraction
ADME or ADMET: Absorption, Distribution, Metabolism, Excretion, and Toxicity
RCSB-PDB: Research Collaboratory for Structural Bioinformatics-Protein Data Bank
AD4: AutoDock4
GA: Genetic Algorithm
LGA: Lamarckian Genetic Algorithm
RMSD: Root Mean Square Deviation
LEU: Leucine
LYS: Lysine
ILE: Isoleucine
PRO: Proline
GLU: Glutamine
TYR: Tyrosine
PHE: Phenylalanine
MET: Methionine
ARG: Arginine
SMILES ID: Simplified Molecular Input Line Entry System identification
GI tract: Gastrointestinal tract
BBB: Blood-Brain Barrier

## Contributions

TS performed the experiment, analysis, graphical designing and wrote the manuscript. AKV led the development of methodology for the experiment, data extraction, study quality assessment, conceptualization, study identification, analysis, manuscript writing and editing, and overall supervision. TS edited the whole draft and did the referencing. NS, NRS and AK provided detailed reviews of the manuscript drafts and delivered crucial critical feedback. The final manuscript, reflecting these collective contributions, has been approved by all authors.

## Declaration of Conflict of Interest

The authors here state that they have no known financial conflicts of interest or personal connections that could have influenced the work presented in this study.

### Acknowledgement

The authors sincerely acknowledge all kinds of support from School of Bioengineering and Biosciences, Lovely Professional University, Punjab, India. Also, the authors also extend their heartfelt thanks to SCFBio, Indian Institute of Technology (IIT)-Delhi, Special Centre for Nanoscience, Jawaharlal Nehru University (JNU), New Delhi, and Gene Regulatory Laboratory, National Institute of Immunology (NII), New Delhi, India for providing computational facilities to conduct our study.

## Funding Declaration

No funding was required for this work.

## Notes

### Competing Interest Statement

The authors have declared no competing interest.

### Summary of Updates

Minor changes have done regarding content of the paper.

