## Supplementary_files_OP_HSA for "*In Silico* Toxicity Assessment of Organophosphates: A DFT and Molecular Docking Study on Their Interaction with Human Serum Albumin (HSA)"

**Supplementary Figure S1:** Graphical abstract

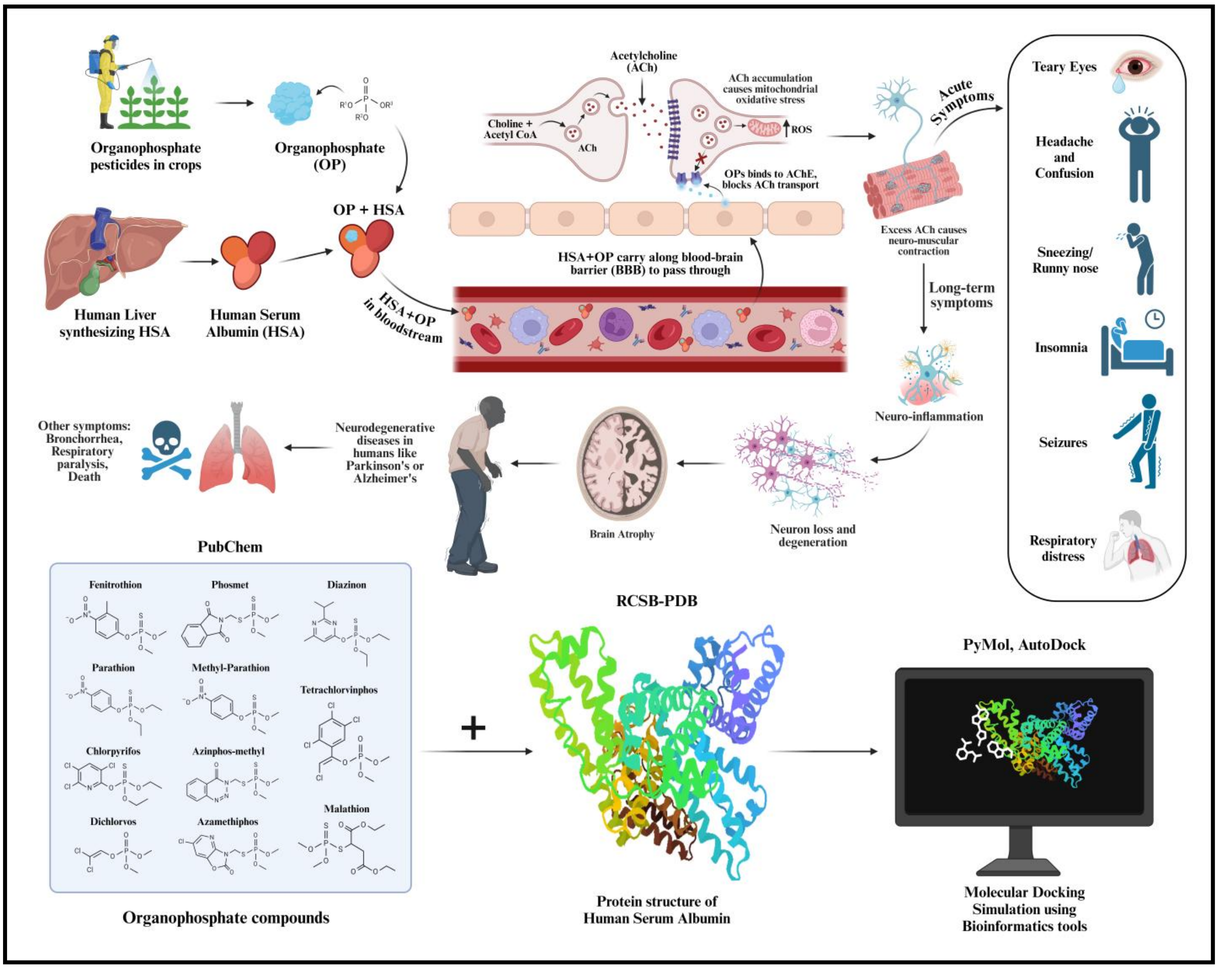


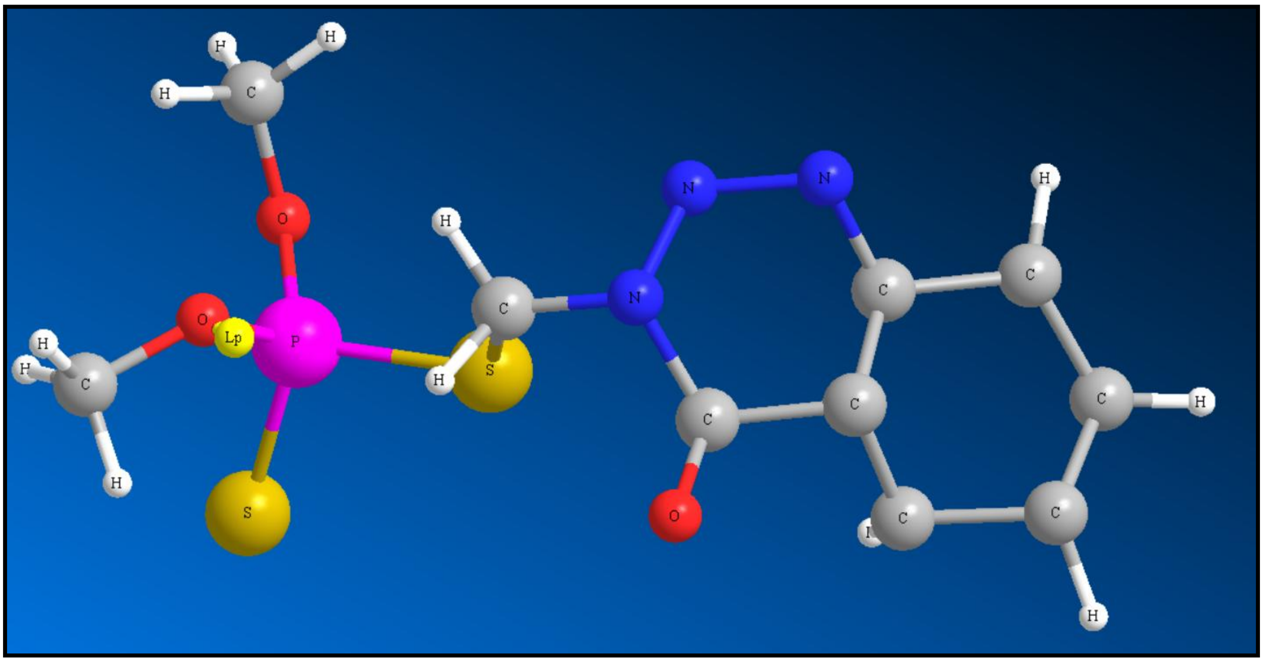

**Supplementary Figure S2:** 3D ground state configuration generated for Azinphos-methyl compound using ChemDraw software.

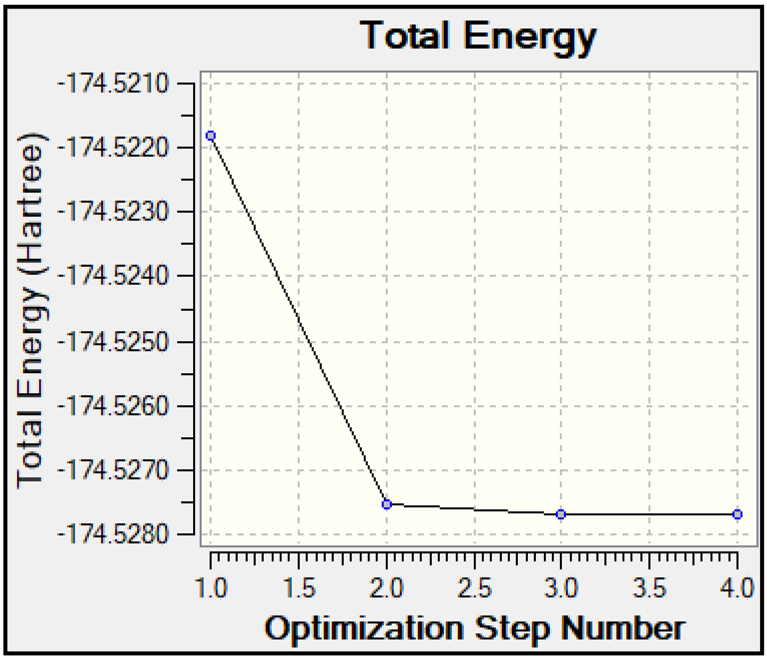


**Supplementary Figure S3:** Graph showing energy minimization for the OP compound Azinphos-methyl using Gaussian09 software.

Internal Coordinates

| | Atom | Bond Atom | Bond Length (Å) | Angle Atom | Angle (°) | 2nd | 2nd Angle (°) | 2nd Angle Type |
|---|---|---|---|---|---|---|---|---|
| 1 | S(1) | | | | | | | |
| 2 | P(3) | S(1) | 2.1267 | | | | | |
| 3 | O(4) | P(3) | 1.6348 | S(1) | 106.8838 | | | |
| 4 | O(5) | P(3) | 1.6354 | S(1) | 107.0049 | O(4) | 97.9348 | Pro-R |
| 5 | S(2) | P(3) | 1.9625 | S(1) | 118.6063 | O(4) | 112.2588 | Pro-S |
| 6 | C(11) | S(1) | 1.8042 | P(3) | 105.3120 | S(2) | -171.9454 | Dihedral |
| 7 | N(7) | C(11) | 1.4523 | S(1) | 110.6933 | P(3) | 179.9797 | Dihedral |
| 8 | H(20) | C(11) | 1.0940 | S(1) | 108.7684 | N(7) | 109.3232 | Pro-R |
| 9 | H(21) | C(11) | 1.0945 | S(1) | 110.0604 | N(7) | 109.6310 | Pro-S |
| 10 | N(8) | N(7) | 1.3688 | C(11) | 114.6231 | S(1) | 89.8205 | Dihedral |
| 11 | C(12) | N(7) | 1.3861 | N(8) | 123.8006 | C(11) | 121.5760 | Pro-S |
| 12 | N(9) | N(8) | 1.2496 | N(7) | 122.0698 | C(11) | -179.9803 | Dihedral |
| 13 | C(10) | C(12) | 1.4627 | N(7) | 114.9040 | N(8) | 0.1703 | Dihedral |
| 14 | O(6) | C(12) | 1.2275 | N(7) | 125.4582 | C(10) | 119.6377 | Pro-R |
| 15 | C(13) | N(9) | 1.3995 | N(8) | 119.8051 | N(7) | 0.0172 | Dihedral |
| 16 | C(14) | C(10) | 1.3980 | C(12) | 121.6942 | C(13) | 121.4959 | Pro-R |
| 17 | C(15) | C(13) | 1.3997 | N(9) | 118.9069 | C(10) | 118.4827 | Pro-S |
| 18 | C(16) | C(14) | 1.3928 | C(10) | 119.4394 | C(12) | 179.9596 | Dihedral |
| 19 | H(22) | C(14) | 1.0862 | C(10) | 121.4069 | C(16) | 119.1537 | Pro-S |
| 20 | C(17) | C(15) | 1.3971 | C(13) | 120.6889 | N(9) | -179.9034 | Dihedral |
| 21 | H(23) | C(15) | 1.0873 | C(13) | 120.2159 | C(17) | 119.0952 | Pro-R |
| 22 | H(24) | C(16) | 1.0857 | C(14) | 120.0894 | C(17) | 120.0008 | Pro-S |
| 23 | H(25) | C(17) | 1.0854 | C(15) | 119.9535 | C(16) | 120.0632 | Pro-S |
| 24 | C(18) | O(4) | 1.4181 | P(3) | 120.3961 | S(1) | 69.4345 | Dihedral |
| 25 | C(19) | O(5) | 1.4181 | P(3) | 120.3829 | S(1) | -69.5326 | Dihedral |
| 26 | H(26) | C(18) | 1.0931 | O(4) | 109.6242 | P(3) | 80.6270 | Dihedral |
| 27 | H(27) | C(18) | 1.0924 | O(4) | 111.0475 | H(26) | 109.8436 | Pro-R |
| 28 | H(28) | C(18) | 1.0919 | O(4) | 108.5847 | H(26) | 108.8120 | Pro-S |
| 29 | H(29) | C(19) | 1.0916 | O(5) | 111.1608 | P(3) | 39.3372 | Dihedral |
| 30 | H(30) | C(19) | 1.0933 | O(5) | 109.5975 | H(29) | 109.9128 | Pro-R |
| 31 | H(31) | C(19) | 1.0928 | O(5) | 108.5217 | H(29) | 108.8453 | Pro-S |

**Supplementary Figure S4:** Tabular information about the calculation of bond length, bond angles and the types of bond angle involved in every atom of Azinphos-methyl molecule during energy minimisation and optimization.

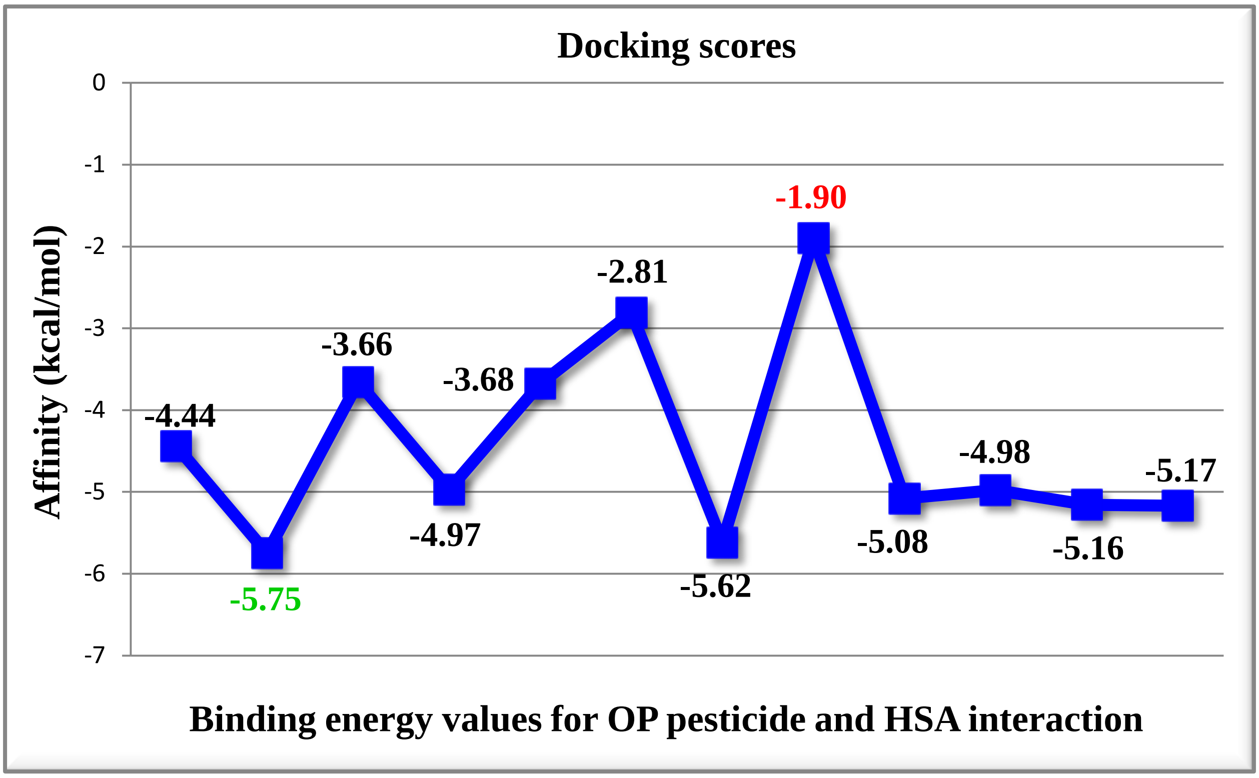


**Supplementary Figure S5:** Graphical plot for docking scores of Organophosphate pesticides (ligand) with HSA (protein). More negative scores indicate more favorable binding affinity.

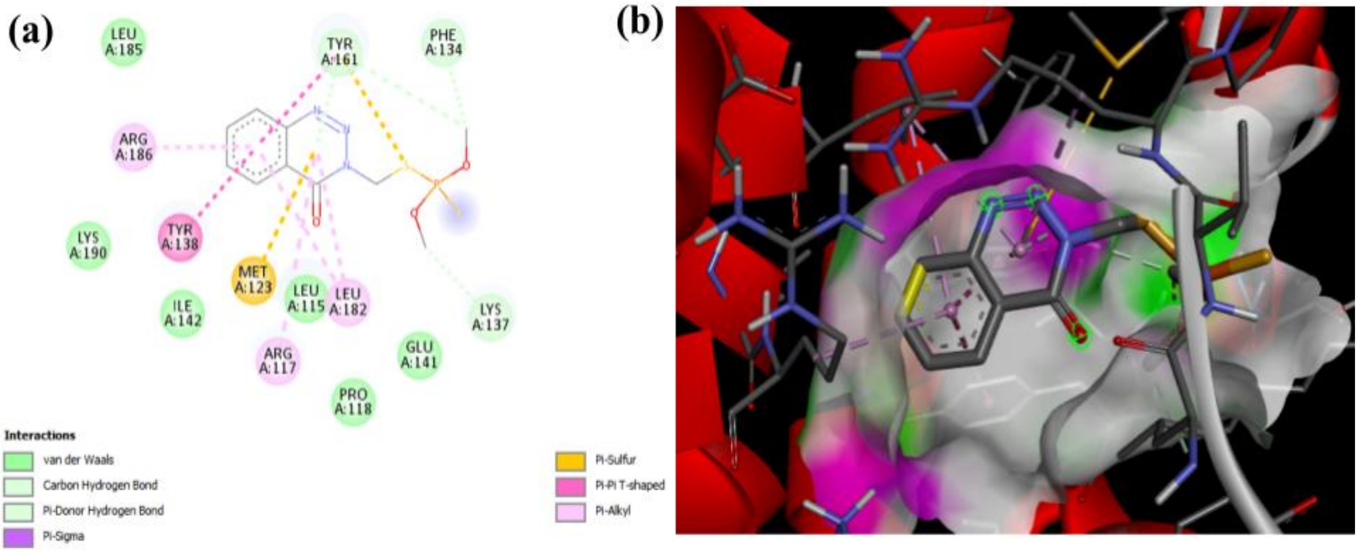

**Supplementary Figure S6:**

(a) 2D representation of non-covalent bonding between Azinphos-methyl and HSA complex;

(b) 3D interaction of Azinphos-methyl with HSA modeled using PyMOL and Discovery Studio software.

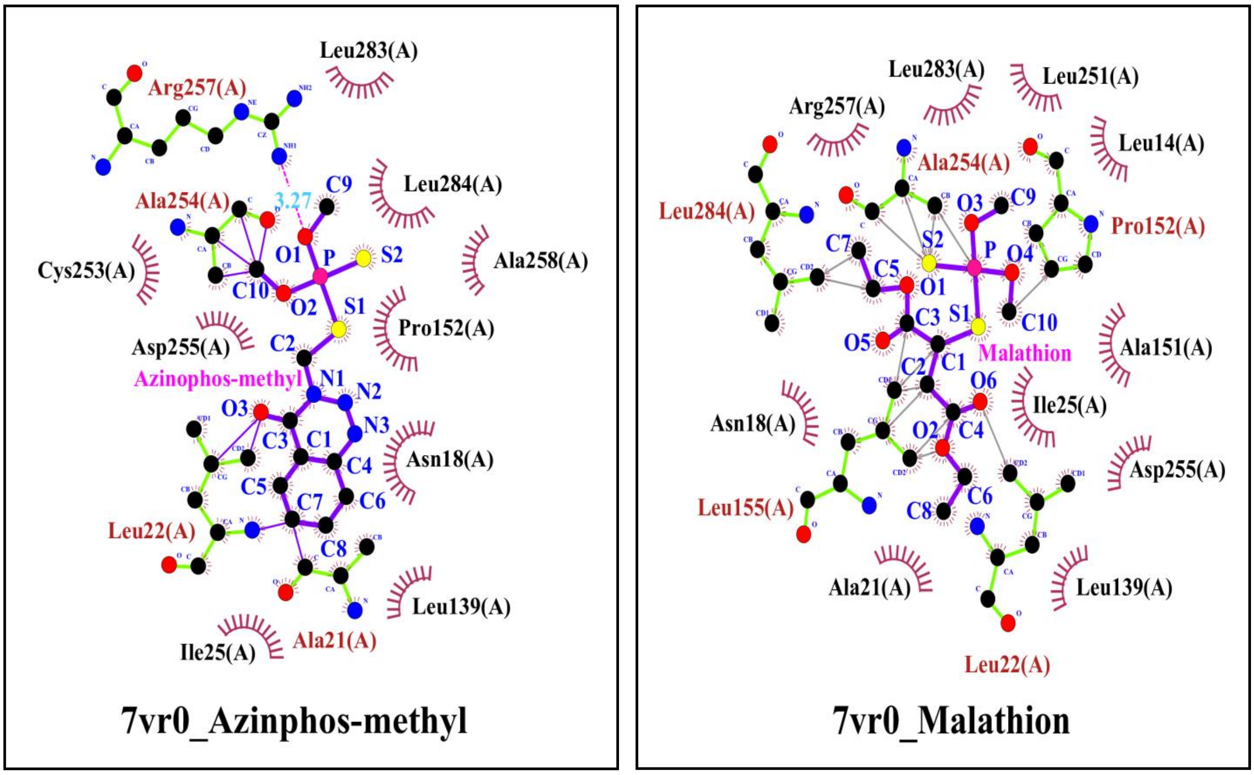


**Supplementary Figure S7:**

(a): Snapshot showing the interaction between HSA and pesticide molecule Azinphos-methyl using LigPlot plus software;

(b): Snapshot showing the interaction between HSA and pesticide molecule Malathion using LigPlot plus software.
